# Impairment of bacteriophage activity in blood: a case study revealing constraints in phage isolation and translation

**DOI:** 10.64898/2026.05.29.728643

**Authors:** Braira Wahid, Teddy Teo, Jinxin Zhao, Lawrence Zang, Anjali Bandara, Qurat-ul-ain Ashraf, Morgyn Warner, Peter Speck

## Abstract

**Background:** Phage therapy is increasingly considered a promising alternative for treating multidrug-resistant (MDR) infections. However, its clinical application remains limited by challenges in isolating effective phages against resistant clinical strains and by the limited ability of in vitro assays to predict performance in real biological environments. While biological matrices are known to influence phage activity, these effects are not well characterised.

**Methods:** A phage-resistant *Pseudomonas aeruginosa* isolate from a patient with recurrent MDR urinary tract infection was used as the model organism. Conventional isolation methods failed to recover effective phages, leading to the development of TEASER-i (Transient EDTA- and Ion-Assisted Sequential Enrichment & Recovery). Recovered phages were characterised using adsorption assays, one-step growth kinetics, and time-kill experiments. Their antibacterial activity was evaluated both *in vitro* and in *ex vivo* human matrices (whole blood, serum, plasma, and urine). Phage efficacy was quantified using maximum log reduction (*E*_*max*_), area under the curve (AUC), and phage-to-bacteria ratio (PBR).

**Results:** A novel TEASER-i method optimised for difficult-to-treat Gram-negative infections, enabled recovery of a functionally effective Øsewage-derived *P. aeruginosa* phage, which outperformed a Øurine-derived *P. aeruginosa* phage that showed slower replication and lower burst size. Phage activity varied significantly in blood, serum, and plasma. Urine supported the most sustained antibacterial effect. In many cases, early bacterial reduction was followed by regrowth. Sustained activity was associated with maintenance of favourable PBR values, while negative PBR corresponded to treatment failure. At 96 h, only two conditions maintained favourable phage load (log _10_ *PBR* > 0): the *S. aureus* phage in urine (+1.66) and the sewage-derived *P. aeruginosa* phage in serum (+1.32).

**Conclusions:** Phage efficacy depends not only on intrinsic lytic capacity but also on the ability to persist and amplify within specific biological environments. Conventional isolation and *in vitro* screening may therefore overestimate therapeutic potential. Combining optimised isolation strategies with *ex vivo* evaluation provides a more realistic framework for phage selection and clinical translation.

## 1. Introduction

The global rise of multidrug-resistant (MDR) bacterial infections has renewed interest in bacteriophages as precision antimicrobials. Despite promising case reports and compassionate-use successes, the clinical translation of phage therapy remains inconsistent, particularly for infections caused by intrinsically phage-resistant or highly adaptive pathogens such as *Pseudomonas aeruginosa* and *Staphylococcus aureus*. A major barrier is the limited ability of current phage discovery pipelines to rapidly identify therapeutically effective phages against difficult-to-treat clinical isolates.

Conventional phage isolation strategies, including direct environmental screening and standard enrichment protocols, are typically optimised for permissive laboratory conditions and assume inherent phage susceptibility [1-5]. However, many clinically relevant bacteria possess multiple defence systems that restrict phage adsorption, replication, or lysis, leading to frequent isolation failure or recovery of weakly lytic phages [6]. Even when plaques are obtained, these often do not translate into promising killing activity *in vivo*, highlighting a critical gap between *in vitro* isolation success and functional therapeutic efficacy [7].

Importantly, most phage characterisation studies rely heavily on *in vitro* assays performed in nutrient-rich media, which fail to recapitulate the physicochemical and immunological complexity of host environments [8]. Biological matrices such as blood, serum, plasma, and urine contain proteins, immune components, and cellular factors that can modulate phage stability, adsorption, and infectivity [9]. As a result, *in vitro* assays frequently overestimate phage killing capacity, providing limited predictive value for clinical performance. This discrepancy has been implicated in several unsuccessful or inconclusive clinical trials, where phages with demonstrated *in vitro* activity failed to achieve therapeutic benefit [10].

*Ex vivo* models offer a critical intermediate between *in vitro* systems and complex *in vivo* as well as clinical settings. These models enable controlled, quantitative assessment of phage-host environment interactions while preserving key physiological constraints that influence phage activity. However, systematic integration of such models into phage discovery and evaluation pipelines remains limited, and their utility for predicting therapeutic success is underexplored. In parallel, there is a growing need for improved strategies to overcome intrinsic phage resistance during isolation. Approaches that transiently modify bacterial surface accessibility or enhance phage adsorption may enable recovery of otherwise undetectable lytic phages, thereby expanding the therapeutic phages specifically against difficult-to-treat bacteria.

Here, we address these challenges by combining an optimised phage isolation strategy, TEASER-i (Transient EDTA- and Ion-Assisted Sequential Enrichment & Recovery), specifically designed for difficult-to-treat clinical isolates, with a quantitative evaluation of phage activity across clinically relevant *ex vivo* matrices. Using both patient-derived and sewage-derived novel phages targeting MDR pathogens, we apply a pharmacodynamic framework to assess antibacterial efficacy, phage persistence, and phage-bacteria dynamics over time in three reference phages: Gram-positive (Ø*S. aureus* phage) and Gram-negative (ØUrine-derived *P. aeruginosa* phage and ØSewage-derived *P. aeruginosa* phage). We hypothesise that (i) conventional isolation approaches underestimate the recoverable phage diversity against resistant pathogens, and (ii) *Ex vivo* matrix-dependent effects are critical in determining whether *in vitro* phage activity results in sustained antibacterial efficacy *ex vivo*. This study establish a baseline that facilitates bridging the gap between laboratory discovery and clinical application.

## 2. Methods

### 2.1. Clinical isolate and urine-derived phage recovery

A patient with a past medical history of recurrent MDR urinary tract infections (UTIs) was presented at Adelaide Hospital. The urine cultures were found to be positive for *P. aeruginosa* isolate and demonstrated resistance to multiple antibiotic courses, resulting in repeated treatment failures. In the absence of effective antimicrobial options, the isolate was submitted for bacteriophage susceptibility testing to the established phage bank of Australia; however, no suitable lytic phages were identified. Therefore, the clinical isolate was subjected to in-house phage isolation and subsequent functional characterisation at the Speck Laboratory, Flinders University. Evidence of phage activity was detected in the patient’s urine samples, as exposure to the host bacterial lawn produced clear halos consistent with lytic phage presence. Individual plaques were isolated, preserved in 20% glycerol at −80°C, and purified for further analysis. This phage was designated as the Ø urine-derived *P. aeruginosa* phage, representing a matched host-phage pair from the same clinical sample. To facilitate comparative analysis of phage behaviour across bacterial cell envelope types, an MDR *S. aureus* isolate was included as a Gram-positive reference. This approach enabled evaluation of differential phage activity between Gram-negative and Gram-positive pathogens under identical experimental conditions. All clinical samples and bacterial isolates were processed in accordance with institutional ethical and biosafety guidelines.

### 2.2. Ethics Statement

This study was conducted in accordance with relevant institutional guidelines and regulations and was approved by the Flinders University Human Research Ethics Committee (Approval No. 9024) and Institutional Biosafety Committee (Approval No. 8892). The patient was under the clinical care and treatment of co-author Dr. Morgyn Warner from February 2024 to August 2025. Verbal informed consent was obtained directly from the patient for participation and publication of relevant clinical information and associated data. Human blood samples were prospectively collected from participants between 03 March 2026 and 17 April 2026, including collection dates of 03 March 2026, 23 March 2026, and 17 April 2026, for subsequent *ex vivo* phage experiments. Written informed consent was obtained from all participants prior to sample collection. All samples and associated data were handled in accordance with approved ethical protocols, and the authors did not have access to information that could identify individual participants during or after data collection.

### 2.3. Introducing TEASER-I (Transient EDTA- and Ion-Assisted Sequential Enrichment & Recovery) for difficult-to-treat bacteria

Phage isolation was initially attempted using the standard *Phage-on-Tap* protocol [1], in which mixture of sewage samples collected from different sites were enriched with the host *P. aeruginosa* isolate under conventional conditions. No phages with consistent lytic activity was recovered. To address this limitation, we developed an optimised enrichment strategy termed TEASER-I. This approach incorporates a staged ion-chelator conditioning step to enhance phage recovery from resistant clinical isolates. Briefly, environmental samples (3 mL sewage) were combined with 7 mL LB broth and 100 µL of mid-log phase host culture (total volume ∼10.1 mL). EDTA was added to a final concentration of 0.05 mM, and the mixture was incubated for 10-15 min at 37°C to transiently increase outer membrane accessibility without compromising host viability. Following this, divalent cations were restored by supplementing the culture with 10 mM CaCl_2_ and 10 mM MgSO_4_ to re-establish conditions favourable for phage adsorption and infection. Enrichment was then continued under standard incubation conditions. Sequential enrichment cycles were performed, consisting of ∼6 h incubation periods followed by transfer into fresh host culture supplemented with divalent cations. This process was repeated daily for 5 consecutive days to selectively amplify rare or weakly infective phage populations. A final overnight amplification step was used to maximise phage titre followed by phage purification. Phage-containing supernatants were subsequently filtered by centrifugation and filtration (0.22 µm), followed by plaque isolation, propagation, and concentration using standard methods. Purified phage stocks obtained through this approach were designated as the Ø sewage-derived *P. aeruginosa* phage

### 2.4. Phage adsorption kinetics

Phage adsorption was quantified by measuring the proportion of unadsorbed phage remaining in the supernatant over time. Briefly, 100 µL of phage suspension was mixed with 100 µL of mid-log phase bacterial culture in 9.8 mL fresh medium. The mixture was incubated at 37°C, and aliquots were collected at 30 s, 1, 2, 4, 6, 8, 10, 15 and 20 min. Samples were centrifuged to pellet bacteria with adsorbed phage, and the supernatant was filtered through a 0.22 µm filter. The filtered supernatant, representing unadsorbed phage, was serially diluted and plated to determine PFU/mL.

The percentage of unadsorbed phage was calculated as:

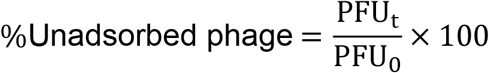

where *PFU*_*t*_ is the titre of unadsorbed phage at each timepoint and *PFU*_0_is the initial phage titre.

Adsorbed phage was then calculated as:

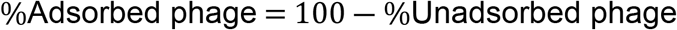

### 2.5. One-step growth assay and time-kill kinetics

One-step growth assay and *in vitro* time-kill kinetics were performed at an MOI of 10, using methods described in our previous studies [5, 11].

### 2.6. *Ex vivo* infection model in human biological matrices

*Ex vivo* phage activity was evaluated using freshly collected human blood obtained from healthy donors (12 mL per donor). Blood samples were processed to generate whole blood, serum, and plasma fractions using standard protocols. All experiments were performed using 1 mL aliquots of each matrix. Bacterial cultures (10^8^ CFU/mL; 20 µL) and corresponding phages (10^9^ PFU/mL; 20µL) were added to each matrix under controlled conditions, and samples were incubated at 37°C in a shaking incubator. At defined timepoints (2, 4, 6, 24, 48, 72, and 96 h), aliquots (10 µL) were collected from each condition, serially diluted, and plated to quantify bacterial load (CFU/mL) and phage titre (PFU/mL) using standard plating methods. Parallel conditions included bacteria-only controls, phage-only controls, and phage-bacteria co-cultures.

### 2.7. Pharmacodynamic analysis of phage activity across biological matrices

Bacterial and phage dynamics were analysed using time-relative CFU/mL and PFU/mL measurements obtained across all matrices and conditions (0-96 h). Phage antibacterial activity was quantified relative to untreated controls using a pharmacodynamic framework [12, 13].

#### Antibacterial effect (E)

The antibacterial effect at each timepoint was calculated as:

*E*(*t*) = log _10_(CFU_control,*t*_) ― log _10_(CFU_treated,*t*_)where *E*(*t*) > 0indicates bacterial reduction in the treated condition relative to the control, and *E*(*t*) ≤ 0indicates no effect or bacterial growth exceeding the control.

From the time-course data, the following summary parameters were derived:

- *E*_max_: maximum observed antibacterial effect
- 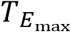: timepoint at which *E*_max_occurred
- *E*_96*h*_: antibacterial effect at 96 h

#### Phage-to-bacteria ratio (PBR)

To quantify phage load relative to bacterial burden, the phage-to-bacteria ratio (PBR) was calculated using matched phage-bacteria co-culture measurements:

log _10_*PBR*_*t*_ = log _10_(PFU_phage+bacteria,*t*_) ― log _10_(CFU_treated,*t*_)where positive values indicate phage abundance exceeding bacterial density, values near zero indicate approximate balance, and negative values indicate bacterial dominance.

### 2.8. Statistical analysis

All experiments were performed in independent triplicates. Data are presented as mean ± standard deviation (SD). Graphs and data visualisation were generated using GraphPad Prism.

## 3. Results

### 3.1. Phage isolation from patient’s urine

A clinical case of recurrent multidrug-resistant urinary tract infection (MDR UTI) was investigated following the failure of multiple antibiotic regimens (**Fig. 1a**). The *P. aeruginosa* isolate was submitted to established phage banks, but no matching lytic phages were identified, resulting in the classification of the strain as phage-resistant. Further attempts using standard isolation methods, including the widely adopted “Phage on Tap” protocol and related phage isolation methods [1], did not yield active phages against the clinical isolate. Direct screening of the patient’s urine on blood agar plates yielded turbid plaques indicating the presence of phage in patient’s urine (named ØUrine-derived PA phage) (**Fig. 1a**). These urine-derived phages demonstrated poor downstream activity, displayed by a decrease in plaque size during purification, slow plaque formation, inefficient amplification, and weak lytic activity, which suggests limited therapeutic potential.

**Figure 1.**
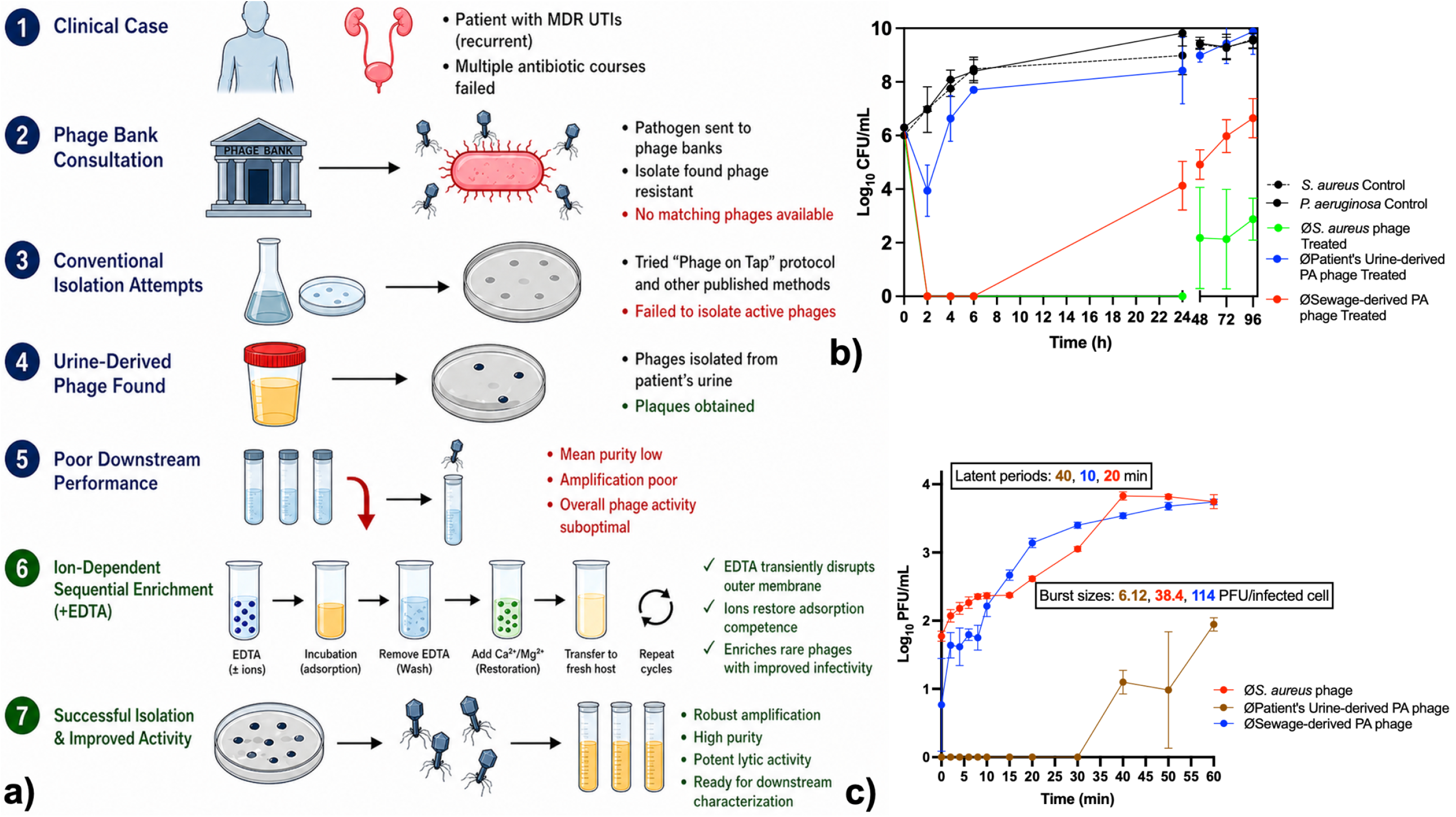
Clinical workflow, phage isolation challenges, and replication kinetics of recovered phages. **(a)** Schematic overview of the clinical case and phage isolation workflow. **(b)** In vitro killing kinetics of *S. aureus* and *P. aeruginosa* in the presence of different phages Gram-positive (Ø*S. aureus* phage) and Gram-negative (ØUrine-derived *P. aeruginosa* phage and ØSewage-derived *P. aeruginosa* phage). **(c)** One-step growth analysis showing replication kinetics of phages. The sewage-derived *P. aeruginosa* phage displayed a short latent period and high burst size, whereas the urine-derived phage showed delayed replication and low burst size.

### 3.2. TEASER-I: a novel optimisable phage isolation method for “difficult-to-treat” bacteria

To overcome these limitations, we developed an ion-dependent sequential enrichment strategy incorporating transient EDTA treatment followed by restoration with divalent cations (Ca^2+^/Mg^2+^) (**Fig. 1a**). This approach was designed to modulate bacterial surface accessibility and enhance phage adsorption. Iterative enrichment cycles led to the recovery of phage ØSewage-derived PA phage with significantly improved amplification efficiency, purity, and lytic activity.

### 3.3. *In vitro* killing dynamics reveal major differences in phage fitness

We next compared the *in vitro* antibacterial activity of ØUrine-derived PA phage and ØSewage-derived PA phage, alongside a *S. aureus* phage as a Gram-positive reference (**Fig. 1b**).

In untreated controls, both *P. aeruginosa* and *S. aureus* exhibited rapid growth, reaching ∼9-10 log_10_ CFU/mL by 24-48 h. In treated bacteria, ØSewage-derived PA phage induced immediate and sustained bacterial suppression, with complete clearance observed as early as 2 h (0 log_10_ CFU/mL), and maintained for up to 6 h. Although partial regrowth occurred at later timepoints (24-96 h), bacterial densities remained substantially reduced (∼3-7 log_10_ CFU/mL) compared to untreated controls, indicating strong initial lytic activity with partial long-term suppression of bacterial growth. In contrast, the ØUrine-derived PA phage, despite being isolated from the same infection context, showed significantly inferior performance. Only a transient reduction in bacterial load was observed at early timepoints (2 h), followed by rapid recovery to control-level bacterial burden (∼9-10 log_10_ CFU/mL by 24 h), demonstrating weak killing and poor replication kinetics. For comparison with Gram-positives, the *S. aureus* phage exhibited rapid early killing, achieving complete bacterial clearance at 2-6 h. However, similar to the sewage-derived *P. aeruginosa* phage, regrowth was observed at later timepoints (48-96 h), suggesting the emergence of resistant subpopulations during late timepoints.

One-step growth analysis showed clear differences in how the phages replicate. The ØSewage-derived PA phage infected quickly, with a short latent period of about 10 minutes and a high burst size of around 114 PFU per infected cell. The ØUrine-derived PA phage took longer to replicate, with a latent period of about 40 minutes and a much lower burst size of about 6 PFU per infected cell. The *S. aureus* phage had intermediate results, with a latent period of about 20 minutes and a burst size of about 38 PFU per infected cell, which matches its moderate antibacterial effect.

### 3.3. Biological matrices differentially reshape phage killing activity

Phage activity was evaluated in whole blood, serum, plasma, and urine (**Fig. 2**). Although all phage stocks were prepared at high titre (∼10^9^ PFU/mL), a rapid reduction in detectable activity was observed at 0 h in all matrices, specifically blood, serum, and plasma.

**Figure 2.**
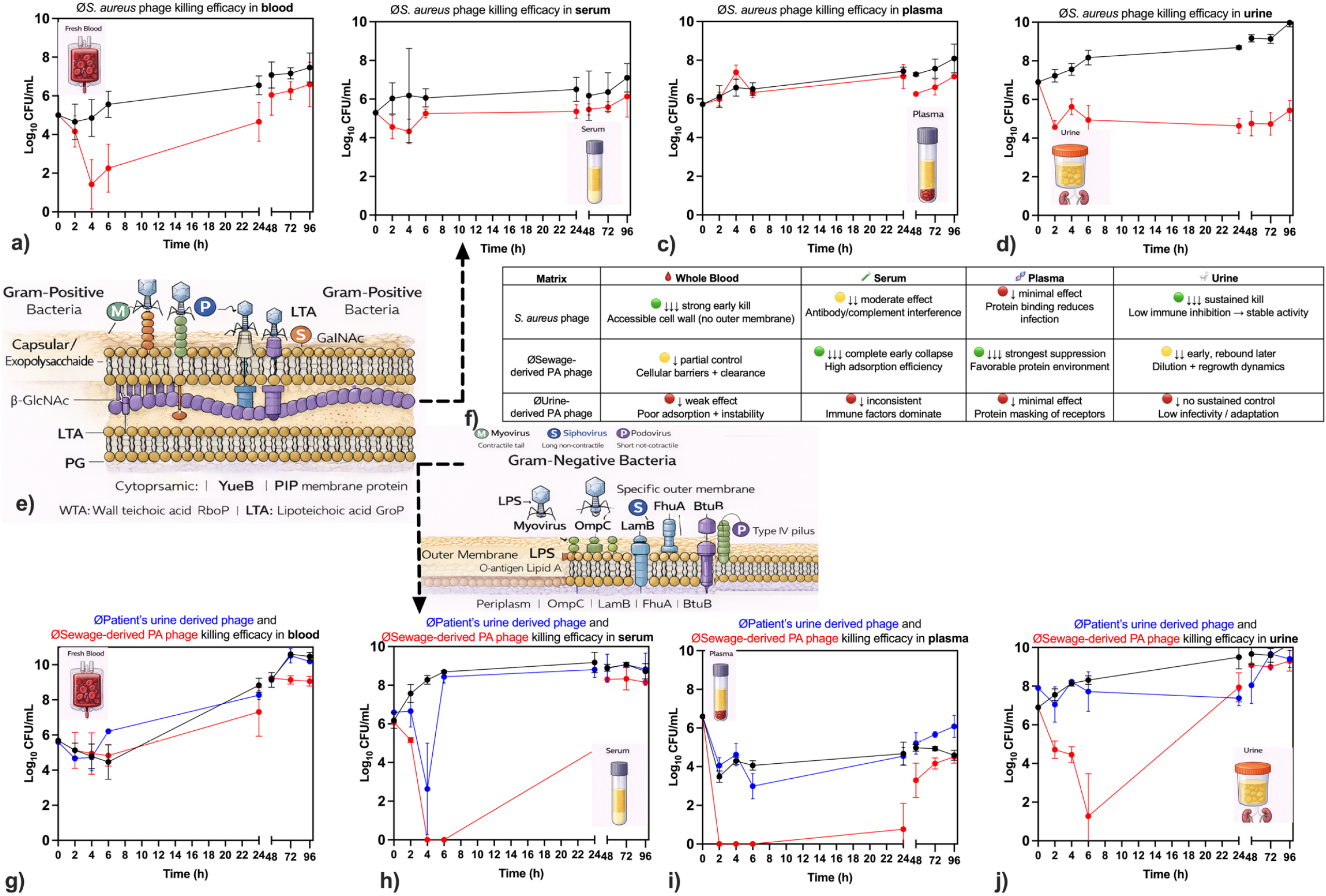
Matrix-dependent variation in phage antibacterial activity and proposed mechanistic framework. **(a-d)** Time-kill curves showing antibacterial activity of the *S. aureus* phage across biological matrices: whole blood **(a)**, serum **(b)**, plasma **(c)**, and urine **(d).** Strong and sustained killing was observed in urine, whereas blood-derived matrices showed transient or reduced activity. **(g-j)** Comparative killing profiles of *P. aeruginosa* phages (sewage-derived and patient urine-derived) across whole blood **(g)**, serum **(h)**, plasma **(i)**, and urine **(j)**. The sewage-derived phage exhibited rapid early killing followed by regrowth, while the urine-derived phage showed weak or inconsistent activity. **(e)** Schematic representation of Gram-positive bacterial cell envelope architecture, highlighting accessibility of phage receptors in the absence of an outer membrane. **(f)** Comparative summary of matrix-dependent effects on phage activity across Gram-positive and Gram-negative systems, integrating observed killing patterns with potential biological constraints.

Matrix-dependent differences observed in *S. aureus phage* are shown in **Fig. 2a-d**. Whole blood supported transient early suppression, with maximal effect at 4 h, followed by progressive regrowth such that treated bacterial counts approached those of the control by 96 h (**Fig. 2a**). Serum produced intermediate and sustained suppression, whereas plasma showed minimal activity overall (Fig. 2b-c). In contrast, urine provided the maximum and sustained antibacterial effect, maintaining substantial suppression through 96 h (Fig. 2d). These findings align with differences in matrix composition: urine likely presents a less inhibitory environment for phage activity, while plasma and blood components may partially impede infection.

Differential profile of both *P. aeruginosa* phages, ØSewage-derived PA phage and ØUrine-derived PA phage, is demonstrated in **Fig. 2g-j**. The ØSewage-derived PA phage exhibited rapid and significant early bacterial killing in serum, plasma, and urine, achieving near-complete suppression at 2-6 h. However, this effect was not sustained, as bacterial regrowth occurred by 24-96 h (**Fig. 2h-j**). In whole blood, the phage displayed only partial and delayed control (**Fig. 2g**). In contrast, the ØUrine-derived PA phage demonstrated weak and inconsistent activity across all matrices throughout the course of infection (**Fig. 2g-j**), consistent with its limited replication kinetics shown in **Fig. 1b and c**.

A comparative summary of these trends is presented in **Fig. 2f**, emphasising matrix- and phage-specific effects. Although the underlying mechanisms were not directly investigated, the observed differences correspond with established structural and biochemical factors (**Fig. 2e-f**). For Gram-positive *S. aureus*, the lack of an outer membrane may enhance phage access to cell wall receptors, especially in less inhibitory environments such as urine. In contrast, Gram-negative *P. aeruginosa* possesses an outer membrane barrier, rendering phage infection more susceptible to matrix-dependent influences on receptor accessibility, adsorption efficiency, or phage stability. Furthermore, serum proteins, complement factors, and plasma-associated molecules may differentially modulate phage-host interactions, leading to reduced or transient activity in blood-derived matrices.

### 3.4. Analysis of phage activity across biological matrices using a pharmacodynamic model

Phage efficacy was quantified using a pharmacodynamic framework based on bacterial reduction relative to untreated controls. The antibacterial effect at each time point was defined as:

*E*(*t*) = log _10_(CFU_control,*t*_) ― log _10_(CFU_treated,*t*_)From this, three summary parameters were derived: the maximum antibacterial effect (*E*_max_), the time to maximum effect 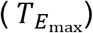, and the residual effect at 96 h (*E*_96*h*_). Across matrices, phage activity was strongly matrix-dependent. The *S. aureus* phage exhibited the most sustained antibacterial activity in urine, where maximal and final effects were comparable, indicating sustained bacterial suppression till late timepoints. In contrast, in whole blood and serum, early killing was observed but was not sustained over time. Specifically, maximal effects occurred at 4 h (*E*_max_ = *E*_4*h*_ ≈ 3.43 in whole blood and 1.86 in serum), followed by progressive decline, with limited residual activity at 96 h (*E*_96*h*_ ≈ 0.87–0.96). Activity in plasma was weaker overall, with delayed and intermediate-to-weak killing effects.

The two Gram-negative *P. aeruginosa* phages displayed significantly different killing profiles. The ØSewage-derived PA phage produced the strongest early antibacterial activity, particularly in serum and urine, with maximal effects at 6 h (*E*_max_ ≈ 8.69 and 7.05, respectively). However, this effect was not sustained, with only intermediate killing activity at 96 h (*E*_96*h*_ ≤ 0.88), indicating substantial regrowth. In contrast, the ØUrine-derived PA phage showed very weak or no activity across all matrices. Although transient early suppression was observed in serum (*E*_max_ ≈ 5.65 at 4 h), this effect was not maintained, and in some matrices (e.g., plasma), negative values of *E*_*96h*_ were observed, indicating bacterial growth exceeding that of the untreated control. Phage persistence varied strongly by matrix and phage type (**Fig. 3**). The *S. aureus* phage remained relatively stable in urine (**Fig. 3d**) but declined or became more variable in blood-derived matrices, particularly serum and plasma (**Fig. 3b-c**), suggesting reduced recoverable phage in protein-rich environments. The ØSewage-derived PA phage showed improved persistence in the presence of bacteria in whole blood and serum (**Fig. 3f-g**), consistent with host-dependent amplification or preservation, whereas plasma reduced its recovery. In contrast, the ØUrine-derived PA phage exhibited poor persistence across most matrices and was rapidly lost under phage-bacteria conditions, particularly in urine (**Fig. 3f-i**). Overall, these findings indicate that sustained phage activity depends not only on intrinsic phage properties but also on whether the surrounding matrix supports recoverable phage survival and amplification.

**Figure 3.**
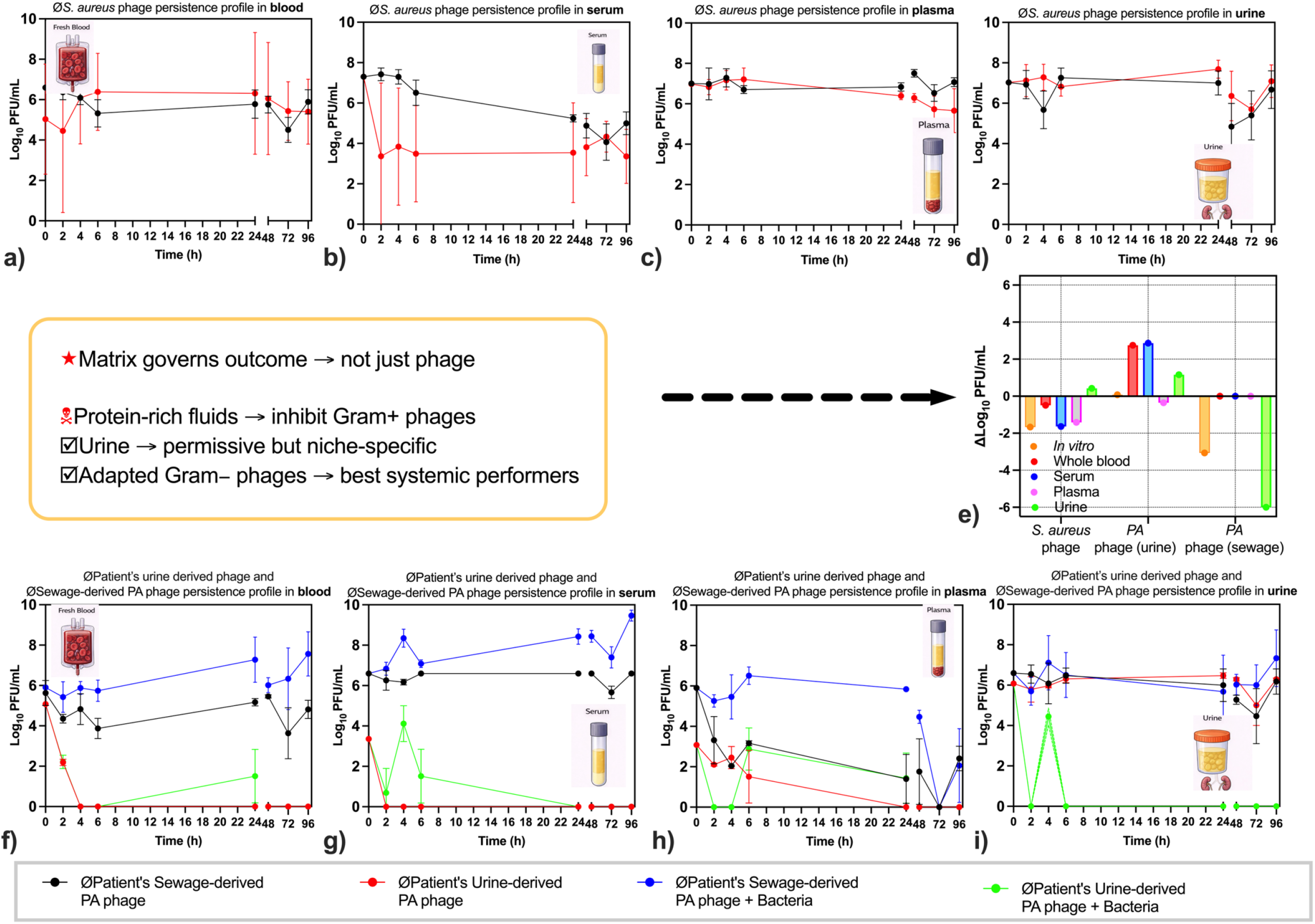
Matrix-dependent phage persistence and host-associated amplification dynamics. **(a-d)** Persistence profiles of the *S. aureus* phage in whole blood **(a)**, serum **(b)**, plasma **(c)**, and urine **(d)**, measured as recoverable PFU/mL over time. Phage stability was highest in urine and more variable in blood-derived matrices. **(f-i)** Persistence of *P. aeruginosa* phages (sewage-derived and urine-derived) in the presence and absence of bacteria across whole blood **(f)**, serum **(g)**, plasma **(h)**, and urine **(i).** The sewage-derived phage showed increased persistence in the presence of bacteria in selected matrices, suggesting host-dependent amplification, whereas the urine-derived phage exhibited rapid loss across conditions. **(e)** Summary of changes in phage recovery (Δlog_10_ PFU/mL) across matrices, comparing phage-only and phage-bacteria conditions.

To pharmacologically interpret these differences, phage-bacteria dynamics were analysed using the phage-to-bacteria ratio (PBR):

log _10_*PBR*_*t*_ = log _10_ (*PFU*_phage+bacteria,*t*_) ― log _10_(*CFU*_treated,*t*_)At 96 h, only two conditions maintained favourable phage load/titre (log _10_*PBR* > 0): the *S. aureus* phage in urine (+1.66) and the sewage-derived *P. aeruginosa* phage in serum (+1.32). All other conditions showed negative PBR values, indicating bacterial dominance. Consistent with this, PFU/mL dataset revealed strong matrix effects. Whole blood and plasma reduced recoverable phage, whereas urine supported higher phage persistence. These exposure differences aligned with antibacterial outcomes, demonstrating that sustained phage availability is required for sustained killing activity. Collectively, these results show that phage efficacy is governed not only by intrinsic lytic activity but by the ability to maintain sufficient phage load/titre within specific biological environments.

## 4. Discussion

This study addresses a central bottleneck in clinical phage therapy: the disconnect between successful phage isolation and reliable therapeutic performance against MDR pathogens. Using a clinically relevant MDR *P. aeruginosa* isolate that failed conventional methods of phage isolation and phage matching with phage banks (**Fig. 1**), we demonstrate that host resistance mechanisms and biological matrix highly constrain both phage recovery and efficacy, and that overcoming these constraints requires both improved isolation strategies and matrix-informed functional characterisation [14].

Failure to identify active phages using established phage banks and standard enrichment approaches highlights a critical limitation of current pipelines, which assume baseline susceptibility in clinical isolates that are difficult to treat and have intrinsic phage defence mechanisms. Even when plaques were recovered directly from patient urine, these phages exhibited poor amplification, low purity, and weak lytic activity, demonstrating that plaque formation alone from biological specimens is not indicative of therapeutic potential.

This observation aligns with prior studies showing that phage isolation success does not necessarily translate into effective bacterial control, particularly in complex biological environments [14, 15], where patient-derived phages often exhibit a narrow host range or limited fitness despite being host-associated [15].

The modified enrichment strategy incorporating transient EDTA exposure followed by Ca^2+^/Mg^2+^ restoration significantly improved phage infectivity and amplification [16]. Mechanistically, this approach likely enhances outer membrane accessibility and adsorption efficiency, consistent with evidence that membrane permeability plays a key role in phage entry and infection kinetics [17]. Interestingly, while higher EDTA concentrations have been shown to directly enhance antibacterial effects when combined with phages [18], our results demonstrate that controlled, low-dose EDTA conditioning can be leveraged during isolation to recover otherwise inaccessible Gram-negative phages without compromising host viability. This represents a conceptual shift: rather than using EDTA as an antimicrobial adjunct, it can be used as a discovery-phase facilitator of phage-host interaction.

Despite being derived from the patient infection site, the urine-derived *P. aeruginosa* phage exhibited significantly inferior killing compared to the sewage-derived phage, both *in vitro* and *ex vivo*. This challenges the assumption that patient-derived phages are inherently optimal for therapy, a concept often promoted in personalised phage approaches and reported in several case studies [14].

Instead, our data show that replication kinetics are the dominant determinant of efficacy, with the sewage-derived phage displaying a short latent period (∼10 min) and high burst size (∼114 PFU/cell), compared to the urine-derived phage (∼40 min latent period; ∼6 PFU/cell). These differences resulted in rapid and significant bacterial clearance during early timepoints, followed by a sustained intermediate killing effect at later timepoints, in ØSewage-derived PA phage (**Fig. 1 b and c**).

Similar relationships between replication efficiency and antibacterial activity have been observed in *ex vivo* systems [19], where phages with rapid killing kinetics achieved early clearance but failed to sustain long-term suppression. Our findings emphasise that phage fitness parameters must be prioritised over source when selecting therapeutic candidates.

A key finding of this study is that biological matrices profoundly reshape phage activity, with substantial differences observed across whole blood, serum, plasma, and urine. While high titres (∼10^9^ PFU/mL) were used, all matrices caused an immediate reduction in recoverable phage at 0 h, indicating rapid loss of infectivity likely due to protein binding, cellular sequestration, or inhibition of adsorption. Consistent with previous *ex vivo* studies, phage efficacy was generally overestimated *in vitro* and attenuated in biologically relevant systems. For example, studies in plant and animal *ex vivo* models report reduced bacterial killing compared to *in vitro* conditions [20], and similar attenuation has been observed in tissue-based systems despite strong initial reductions [21].

Similar to existing literature, our findings also confirm that urine supported the highest phage persistence and most sustained antibacterial effect, and blood, serum, and plasma acted as a restrictive environment for phage propagation and bacterial killing. These findings are consistent with reports that phage stability and distribution vary significantly across biological environments, influencing therapeutic outcomes [22]. Early killing does not predict sustained efficacy. Across all phages, strong early antibacterial effects were frequently followed by bacterial regrowth at later timepoints, particularly in blood-derived matrices. This pattern closely mirrors observations from multiple *ex vivo* and *in vivo* studies, where initial phage-mediated reductions are not sustained without sufficient phage persistence or amplification [22, 23].

Notably, even highly effective phages (e.g., sewage-derived phage) showed declining efficacy over time, indicating that early lytic activity alone is insufficient for long-term control. This highlights the importance of evaluating temporal dynamics rather than endpoint measurements. By integrating PFU and CFU measurements, we show that phage efficacy is governed by the ability to maintain a favourable PBR. Only two conditions maintained positive PBR at 96 h: *S. aureus* phage in urine (+1.66) and Sewage-derived *P. aeruginosa* phage in serum (+1.32) All other conditions showed negative PBR values, indicating bacterial dominance and loss of effective phage load/titre. This explains why even phages with strong early killing ultimately failed to sustain bacterial suppression. This concept is supported by pharmacokinetic/pharmacodynamic *ex vivo* models [19, 21-23], where maintenance of sufficient phage exposure relative to bacterial burden is required to achieve sustained bactericidal activity and prevent resistance emergence.

This study has several limitations that should be considered when interpreting the findings. Experiments were conducted at a relatively small scale, using triplicate measurements and limited matrix volumes (1 mL per condition), which may not fully capture inter-individual variability or the complexity of physiological environments. In addition, while the *ex vivo* system provides a more clinically relevant framework than standard *in vitro* assays, it does not fully recapitulate *in vivo* dynamics such as immune cell regulation, tissue compartmentalisation, and phage pharmacokinetics. These observations, therefore, warrant validation in large-scale experimental systems. Despite these limitations, the findings raise several important questions for the field.

- First, to what extent do current phage isolation pipelines fail to capture functionally relevant phages against highly resistant clinical isolates, and how can enrichment strategies be systematically optimised?
- Second, what are the dominant mechanisms driving phage loss or inactivation in biological matrices, protein binding, immune factors, or physicochemical instability, and how can these be mitigated?
- Third, can quantitative pharmacodynamic parameters, such as phage-to-bacteria ratios and exposure (AUC), be standardised to predict clinical efficacy across different infection contexts?
- Finally, how can *ex vivo* models be further refined to better bridge the gap between laboratory screening and patient outcomes?

Addressing these questions will be critical for advancing phage therapy from experimental promise to a reliable clinical modality, particularly for difficult-to-treat and phage-resistant infections.

## Conclusion

This study demonstrates that phage therapy success is not determined solely by lytic capacity, but by the ability to sustain effective phage load/titre within complex biological environments. By integrating improved isolation with matrix-specific pharmacodynamic evaluation, we provide a framework that more accurately predicts clinical performance and addresses a key gap between laboratory discovery and therapeutic application.

## Ethical approval

All methods were carried out in accordance with relevant institutional guidelines and regulations and were approved by the Flinders University Institutional Biosafety Committee (approval no. 8892) and Human Research Ethics Approval number 9024. The patient was under the clinical care and treatment of co-author Dr. Morgyn Warner from February 2024 to August 2025. Verbal informed consent was obtained directly from the patient for participation and publication of relevant clinical information and associated data. Human blood samples were prospectively collected from participants between 03 March 2026 and 17 April 2026, including collection dates of 03 March 2026, 23 March 2026, and 17 April 2026, for subsequent *ex vivo* phage experiments. Written informed consent was obtained from all participants before sample collection. Environmental water and sewage samples were collected from the Flinders University campus pond (Adelaide, Australia) and publicly accessible wastewater sources associated with SA Water under the same approval and in compliance with local regulations.

## References

[1] N. Bonilla, M. I. Rojas, G. N. F. Cruz, S.-H. Hung, F. Rohwer, and J. J. Barr, “Phage on tap–a quick and efficient protocol for the preparation of bacteriophage laboratory stocks,” PeerJ, vol. 4, p. e2261, 2016.

[2] A. M. Pinto, A. Faustino, L. M. Pastrana, M. Bañobre-López, and S. Sillankorva, “Pseudomonas aeruginosa PAO 1 in vitro time–kill kinetics using single phages and phage formulations—modulating death, adaptation, and resistance,” Antibiotics, vol. 10, no. 7, p. 877, 2021.

[3] F. Dai, G. Yang, J. Lou, X. Zhao, M. Chen, G. Sun, and Y. Yu, “Isolation and characterization of Pseudomonas aeruginosa phages with a broad host spectrum from hospital sewage systems and their therapeutic effect in a mouse model,” The American Journal of Tropical Medicine and Hygiene, vol. 108, no. 6, p. 1220, 2023.

[4] S. Sharma, S. Datta, S. Chatterjee, M. Dutta, J. Samanta, M. G. Vairale, R. Gupta, V. Veer, and S. K. Dwivedi, “Isolation and characterization of a lytic bacteriophage against Pseudomonas aeruginosa,” Scientific Reports, vol. 11, no. 1, p. 19393, 2021.

[5] B. Wahid, S. C. Nang, J. Zhao, Z. Y. Kho, M. Han, H. H.-Y. Yu, H. Wickremasinghe, P. J. Bergen, S. Aslam, and R. T. Schooley, “Phage Resistance Bidirectionally Altered Antibiotic Susceptibility in Klebsiella pneumoniae via galE mutation,” International Journal of Antimicrobial Agents, pp. 107738, 2026.

[6] A. S. Nilsson, “Pharmacological limitations of phage therapy,” Upsala journal of medical sciences, vol. 124, no. 4, p. 218–227, 2019.

[7] J. Lin, F. Du, M. Long, and P. Li, “Limitations of phage therapy and corresponding optimization strategies: a review,” Molecules, vol. 27, no. 6, p. 1857, 2022.

[8] S. A. Gomez-Ochoa, M. Pitton, L. G. Valente, C. D. S. Vesga, J. Largo, A. C. Quiroga-Centeno, J. A. H. Vargas, S. J. Trujillo-Caceres, T. Muka, and D. R. Cameron, “Efficacy of phage therapy in preclinical models of bacterial infection: a systematic review and meta-analysis,” The Lancet Microbe, vol. 3, no. 12, p. e956–e968, 2022.

[9] K. Danis-Wlodarczyk, K. Dąbrowska, and S. T. Abedon, “Phage therapy: the pharmacology of antibacterial viruses,” Current issues in molecular biology, vol. 40, no. 1, p. 81–164, 2021.

[10] J. Bull, B. R. Levin, T. DeRouin, N. Walker, and C. A. Bloch, “Dynamics of success and failure in phage and antibiotic therapy in experimental infections,” BMC microbiology, vol. 2, no. 1, p. 35, 2002.

[11] B. Wahid, J. Zhao, A. Truskewycz, M. MacGregor, J. Ramsay, M. Warner, and P. Speck, “Ionic regulation of Gram-positive phage adsorption governs host range and improves phage isolation efficiency,” Communications Biology, vol. Under-review, 2026.

[12] Y. Lin, R. Y. Chang, G. Rao, B. Jermain, M.-L. Han, J. Zhao, K. Chen, J. Wang, J. Barr, and R. T. Schooley, “Pharmacokinetics/pharmacodynamics of antipseudomonal bacteriophage therapy in rats: a proof-of-concept study,” Clinical Microbiology and Infection, vol. 26, no. 9, p. 1229–1235, 2020.

[13] A. Echterhof, T. Dharmaraj, P. Blankenberg, B. Targ, T. D. Nguyen, P. L. Bollyky, N. M. Smith, and F. G. Blankenberg, “Whole-body distribution of three Pseudomonas phages characterized by a translational physiologically based pharmacokinetic model,” Antimicrobial Agents and Chemotherapy, vol. 70, no. 1, p. e01506–25, 2025.

[14] M. Pitton, L. G. Valente, S. Oberhaensli, B. Gözel, S. M. Jakob, P. Sendi, M. Fürholz, D. R. Cameron, and Y.-A. Que, “Targeting chronic biofilm infections with patient-derived phages: an in vitro and ex vivo proof-of-concept study in patients with left ventricular assist devices.” p. ofaf158.

[15] M. Almosuli, A. Kirtava, A. Chkhotua, L. Tsveniashvili, N. Chanishvili, S. S. Irfan, E. Ng, H. McIntyre, A. J. Hockenberry, and R. P. Araujo, “Urinary bacteriophage cooperation with bacterial pathogens during human urinary tract infections supports lysogenic phage therapy,” Communications Biology, vol. 8, no. 1, p. 175, 2025.

[16] H.-H. Huang, M. Furuta, T. Nasu, M. Hirono, J. Pruet, H. M. Duc, Y. Zhang, Y. Masuda, K.-i. Honjoh, and T. Miyamoto, “Inhibition of phage-resistant bacterial pathogen re-growth with the combined use of bacteriophages and EDTA,” Food microbiology, vol. 100, p. 103853, 2021.

[17] R. Daugelavicius, V. Cvirkaite, A. r. Gaidelytė, E. Bakiene, R. Gabrenaite-Verkhovskaya, and D. H. Bamford, “Penetration of enveloped double-stranded RNA bacteriophages φ13 and φ6 into Pseudomonas syringae cells,” Journal of virology, vol. 79, no. 8, p. 5017–5026, 2005.

[18] L. Imamovic, and M. Muniesa, “Characterizing RecA-independent induction of Shiga toxin2-encoding phages by EDTA treatment,” PloS one, vol. 7, no. 2, p. e32393, 2012.

[19] L. Rivas, B. Coffey, O. McAuliffe, M. J. McDonnell, C. M. Burgess, A. Coffey, R. P. Ross, and G. Duffy, “In vivo and ex vivo evaluations of bacteriophages e11/2 and e4/1c for use in the control of Escherichia coli O157: H7,” Applied and environmental microbiology, vol. 76, no. 21, p. 7210–7216, 2010.

[20] L. A. Pinheiro, C. Pereira, M. E. Barreal, P. P. Gallego, V. M. Balcão, and A. Almeida, “Use of phage ϕ6 to inactivate Pseudomonas syringae pv. actinidiae in kiwifruit plants: In vitro and ex vivo experiments,” Applied microbiology and biotechnology, vol. 104, no. 3, p. 1319–1330, 2020.

[21] A. Dalponte, V. Filor, A. Nerlich, M. Müsken, M. Fulde, and W. Bäumer, “Evaluation of bacteriophage efficacy against Pseudomonas aeruginosa in ex vivo and in vitro canine skin systems,” Scientific reports, 2026.

[22] M. Brack, M. Vollgraf, G. Nouailles, I. Korf, S. Wienecke, A. Dannheim, H. Ziehr, J. Bushe, A. Voss, and A. Gruber, “Phage4Cure–ex vivo and in vivo efficacy testing of a phage cocktail against Pseudomonas aeruginosa,” Pneumologie, vol. 76, no. S 01, p. FV390, 2022.

[23] A. J. Kunz Coyne, K. Stamper, A. El Ghali, R. Kebriaei, B. Biswas, M. Wilson,M. V. Deschenes, T. T. Tran, C. A. Arias, and M. J. Rybak, “Phage-antibiotic cocktail rescues daptomycin and phage susceptibility against daptomycin-nonsusceptible Enterococcus faecium in a simulated endocardial vegetation ex vivo model,” Microbiology Spectrum, vol. 11, no. 4, p. e00340–23, 2023.

